# Neurological disorder drug discovery from gene expression with tensor decomposition

**DOI:** 10.1101/704163

**Authors:** Y-h. Taguchi, Turki Turki

## Abstract

**Background:** Identifying effective candidate drug compounds in patients with neurological disorders based on gene expression data is of great importance to the neurology field. By identifying effective candidate drugs to a given neurological disorder, neurologists would (1) reduce the time searching for effective treatments; and (2) gain additional useful information that leads to a better treatment outcome. Although there are many strategies to screen drug candidate in pre-clinical stage, it is not easy to check if candidate drug compounds can be also effective to human.

**Objective:** We tried to propose a strategy to screen genes whose expression is altered in model animal experiments to be compared with gene expressed differentically with drug treatment to human cell lines.

**Methods:** Recently proposed tensor decomposition (TD) based unsupervised feature extraction (FE) is applied to single cell (sc) RNA-seq experiments of Alzheimer’s disease model animal mouse brain.

**Results:** Four hundreds and one genes are screened as those differentially expressed during A*β* accumulation as age progresses. These genes are significantly overlapped with those expressed differentially with the known drug treatments for three independent data sets: LINCS, DrugMatrix and GEO.

**Conclusion:** Our strategy, application of TD based unsupervised FE, is useful one to screen drug candidate compounds using scRNA-seq data set.

## 1 Introduction

Drug discovery for neurological disorder has never been successful in spite of massive efforts spent [1]. One possible reason is because we generally do not have suitable model animals for human neurological disorder [2]. Although a huge number of compounds are screened using model animals, only a few of them passed the human level screening. In this sense, it is required to screen candidate compounds using information retrieved from human at the earliest stage. One possible strategy to do this is the usage of human cell lines; Nevertheless, it is also not easy to perform, since generating cell line from human neurological disorder patients is not easy. In contrast to the cancer cell lines, which can be easily generated by immortalizing tumor cells, neuronal cells are hardly converted to cell lines, since mature neurons do not undergo cell division [3]. Therefore, it is difficult to test if candidate drugs work for human during pre-clinical stages.

In order to overcome this difficulty, we proposed an alternative strategy; comparing disease gene expression with that of compound treated animals and/or human cell lines. Generally, compound screening is based upon phenotype; i.e., evaluation of compounds efficiency is tested based upon if drug treatment can produce symptomatic improvement. Nevertheless, since it has been recently found that various neurological disorders share gene expression [4], focusing on gene expression profiles might be more reasonable. Following this strategy, we considered gene expression profiles (single cell RNA-seq) of mouse brain during amyloid *β* accumulation. As being aged, some set of gene expression progresses and significantly overlaps with genes that express differential expression caused by various compounds treatment. Since top ranked (i.e., with the most overlaps) detected compounds turn out to be tested previously toward Alzheimer disease (AD) treatment, lower ranked compounds also might be promising candidate compounds for AD.

Expression levels exhibit variations of scRNA-seq data used in this study due to contributions specific to genotypes, tissues, ages, sex, plates, wells, and interactions thereof. Hence, classical unsupervised decomposition methods are not well-suited to explore the six-way interactions and struggle to extract insights from data, hindering the process of finding effective drug compounds of a neurological disorder.

### Contributions

Our contributions over existing work are summarized as follows:

– Whilst the application of tensor decomposition (TD) to the neurology domain is not new, previous developments, to the best of our knowledge, facilitated the neurological drug discovery process are not relevant to modeling the several interactions of scRNA-seq data used in this work. Our proposed tensor decomposition formalism is new, targeting neurological drug discovery of AD and constitutes a main contribution of this work.
– We present findings on an AD with a tensor decomposition formalism demonstrating the effectiveness of finding compounds for the treatment of AD.
– As similar to tensor decomposition techniques, the utilized tensor decomposition technique works under the unsupervised learning setting which is more time effective than previous deployments that work under different learning settings, including the supervised learning setting.
– Unlike traditional machine and deep learning approaches that provide solutions to artificial intelligence when applied to plents of neurological disorder problems, our approach blends techniques from linear algebra and statistics to yield a tensor decomposition technique utilizing a statistical linear algebra approach, requiring much less computational resources and time to reach a solution [5–7].

### Organization

The rest of the paper is organized as follows. Section 2 introduces the tensor decomposition technique and the provided data to be analyzed. Section 3 presents the experimental results, followed by Section 4 to discuss the results. Section 5 concludes the work and points out future direction.

## 2 Materials and Methods

### 2.1 Single cell RNA-seq

Single cell (sc) RNA-seq used in this study was downloaded from gene expression omnibus (GEO) using GEO ID GSE127892. It is composed of two genotypes (APP_NL-F-G and C57Bl/6), two tissues (Cortex and Hippocampus), four ages (3, 6, 12, and 21 weeks), two sex (male and female) and four 96 well plates. For each of combined combinations, four 96 well plates, each of wells includes one cell, were tested. Among those wells tested, wells with insufficient gene expression were discarded. As a result, among 2 (genotype) × 2 (tissues) × 4 (ages) × 2 (sex) × 4 (plates) × 96 (wells) = 12288 cells measured, scRNA-seq for only 10801 cells were provided.

### 2.2 Tensor decomposition based unsupervised feature extraction

We applied recently proposed TD based unsupervised feature extraction (FE) [8 – 18] to scRNA-seq. A tensor 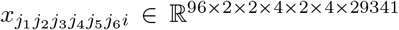 that represents gene expression of *i*th gene of *j*_1_th cell (well) at *j*_2_th genotyoe (*j*_2_ = 1:APP_NL-F-G and *j*_2_ = 2: C57Bl/6), *j*_3_th tissue (*j*_3_ = 1:Cortex and *j*_3_ = 2:Hippocampus), *j*_4_th age (*j*_4_ = 1: three weeks, *j*_4_ = 2: six weeks, *j*_4_ = 3: twelve weeks, and *j*_4_ = 4: twenty one weeks), *j*_5_th sex (*j*_5_ = 1:female and *j*_5_ = 2:male) and *j*_6_th plate.

*x*_*j*_1_*j*_2_*j*_3_*j*_4_*j*_5_*j*_6_*i*_ is standardized such that 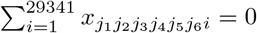 and 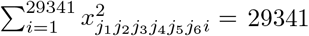. HOSVD [9] was applied to *x*_*j*_1_*j*_2_*j*_3_*j*_4_*j*_5_*j*_6_*i*_ such that

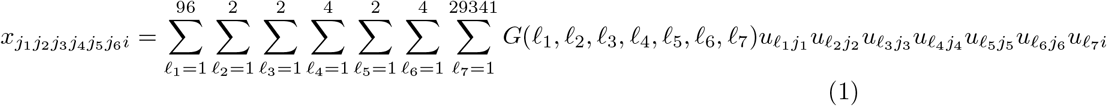

where 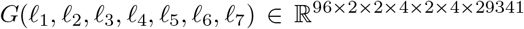 is core tensor, 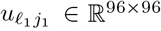, 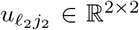, 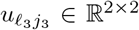, 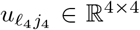, 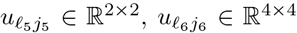 and 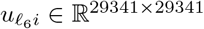 are singular value matrices that are orthogonal matrices. In order to save time to compute, only 1 ≤ *ℓ*_1_, *ℓ*_7_ ≤ 10 were computed (The reason why we employed specifically HOSVD in this research will be discussed in the discussion section, because it is difficult to explain the reason before demonstrating how we make use of TD for data analysis).

After investigation of *u*_*ℓ*_4_*j*_4__, *u*_2_*j*_4___ represent monotonic dependence upon age while *ℓ*_1_, *ℓ*_2_, *ℓ*_3_, *ℓ*_5_, *ℓ*_6_ = 1 represent independence of cells, genotype, tissue, sex and plate. Since *G*(1, 1, 1, 2, 1, 1, 2) has the largest absolute vales among *G*(1, 1, 1, 2, 1, 1, *ℓ*_7_), *u*_2*i*_ is employed to compute *P*-values attributed to *i*th gene as

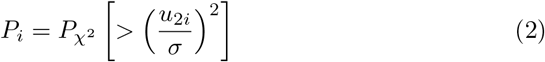

where *P*_*χ*^2^_ [> *x*] is the cumulative probability of *χ*^2^ distribution when the argument is larger than *x* and *σ* is the standard deviation.

*P*-values are corrcted by Benjamini and Hochberg criterion [19] and genes associated with corrected *P*-values less than 0.01 are selected for downstream analysis.

### 2.3 Enrichment analysis

Four hundreds and one genes selected by TD based unsupervised FE were uploaded to Enrichr [20] for enrichment analysis. Full list of enrichment analysis as well as list of 401 genes are accessible at https://amp.pharm.mssm.edu/Enrichr3/enrich?dataset=5bbbe5602715daf9787895cd16829707

List of 401 genes and three enrichment analyses used in this study, “LINCS L1000 Chem Pert up”, “DrugMatrx” and “Drug Perturbations from GEO up” are also available as supplementary material.

Ranks are based upon adjusted P-values (not those provided by Enrichr).

## 3 Results

As a unsupervised technique applied to scRNA-seq data set, we employ tensor decomposition [21] that was sometimes applied to gene expression analysis [22].

### 3.1 Synthetic study of TDs

Before performing TD based unsupervised FE, we perform some synthetic study for some TDs.

We prepared two synthetic data sets, 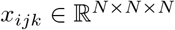 defined as

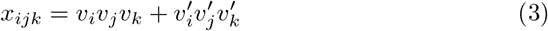

where 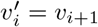 for *i* ≤ *N* – 1 and 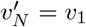.

For data set 1 (Fig. 1(A) and (B)),

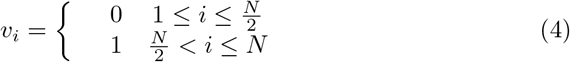

and for data set 2 (Fig. 1(C) and (D)).

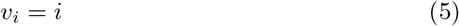

**Figure 1:**
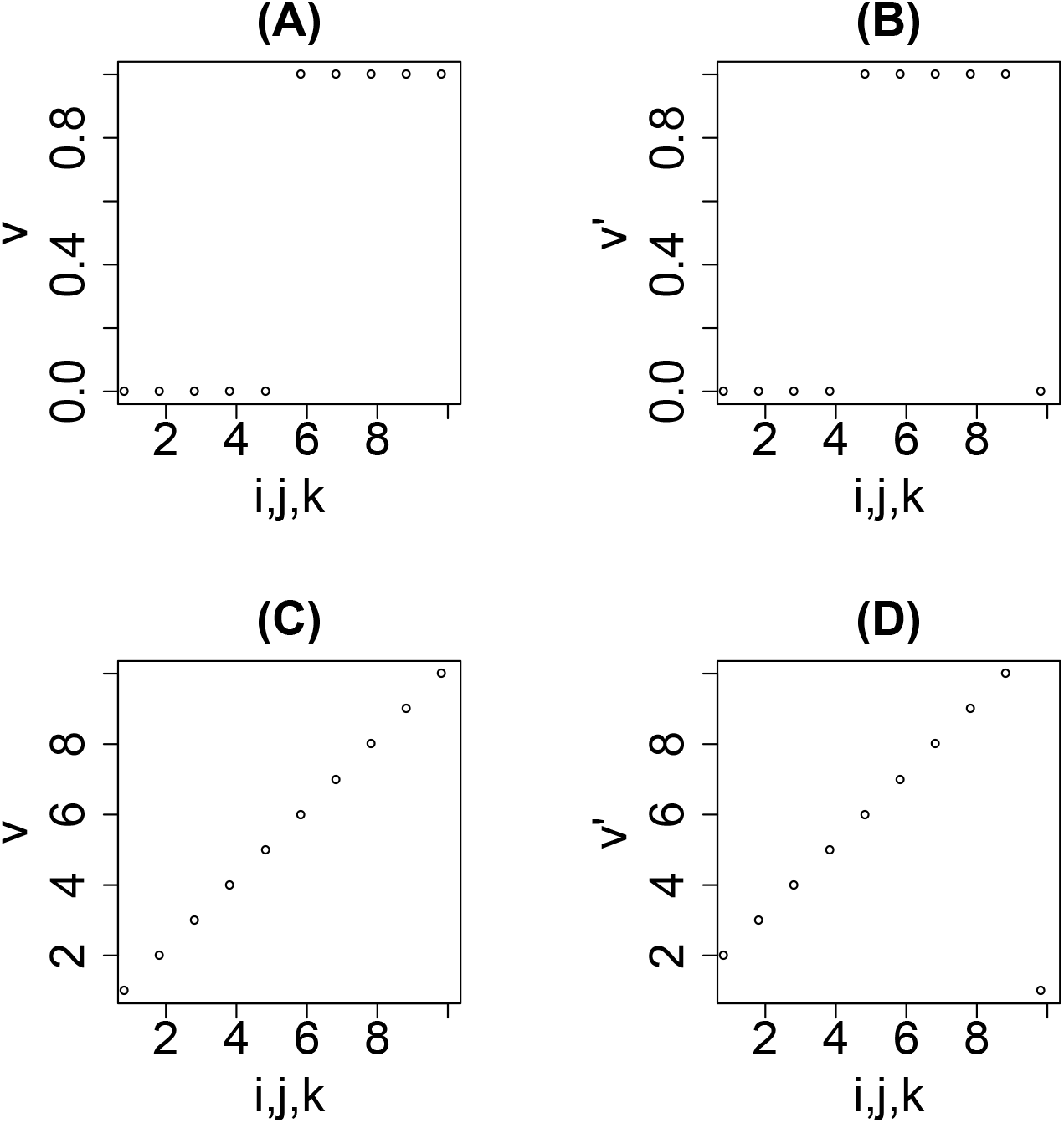
Data set 1, eq. (4), (A) *v_i_* and (B) 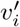 and data set 2, eq. (5), (C) *v_i_* and (D) 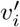.

We apply HOSVD, CP decomposition and CMTF [23] to these two synthetic data set with *N* = 10. At first, we applied HOSVD to data set 1 and 2 as

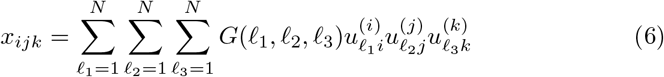

where 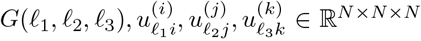. Then we noticed that only four *G*s with (*ℓ*_1_, *ℓ*_2_, *ℓ*_3_) = (1, 1, 1), (1, 2, 2), (2, 1, 2), (2, 2, 1) have non zero values for both data set 1 and 2. Figs. 2 and 3 show 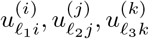 and

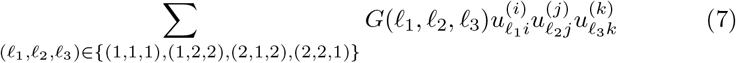

**Figure 2:**
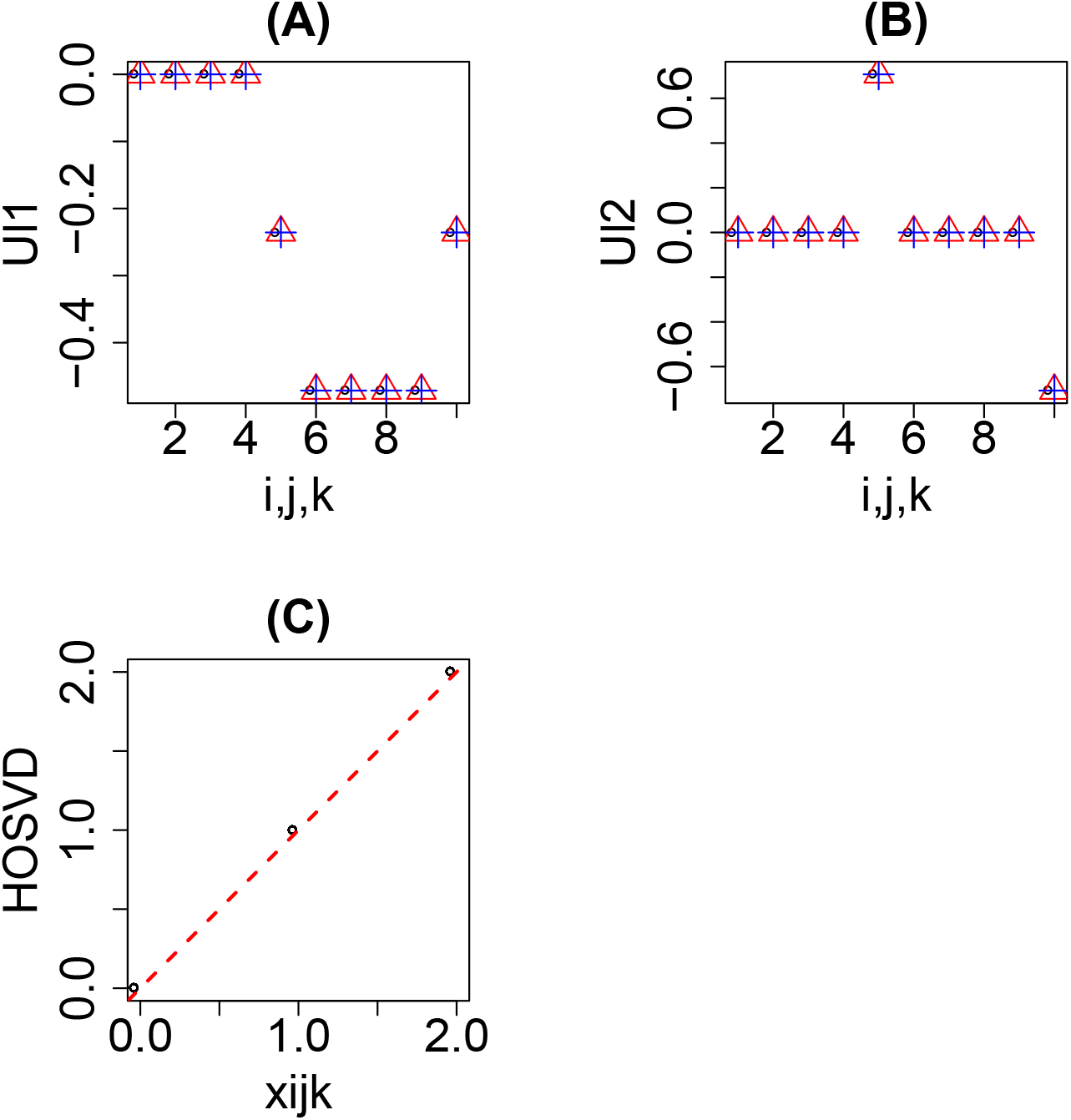
The results obtained by HOSVD applied to data set 1: eq. (4). (A) Open black circles: 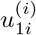, open red triangles: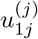, blue pluses: 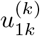 (B) Open black circles: 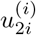, open red triangles: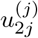, blue pluses: 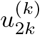. (C) Scatter plot between *x_ijk_* (horizontal axis) and eq. (7) (vertical axis).

**Figure 3:**
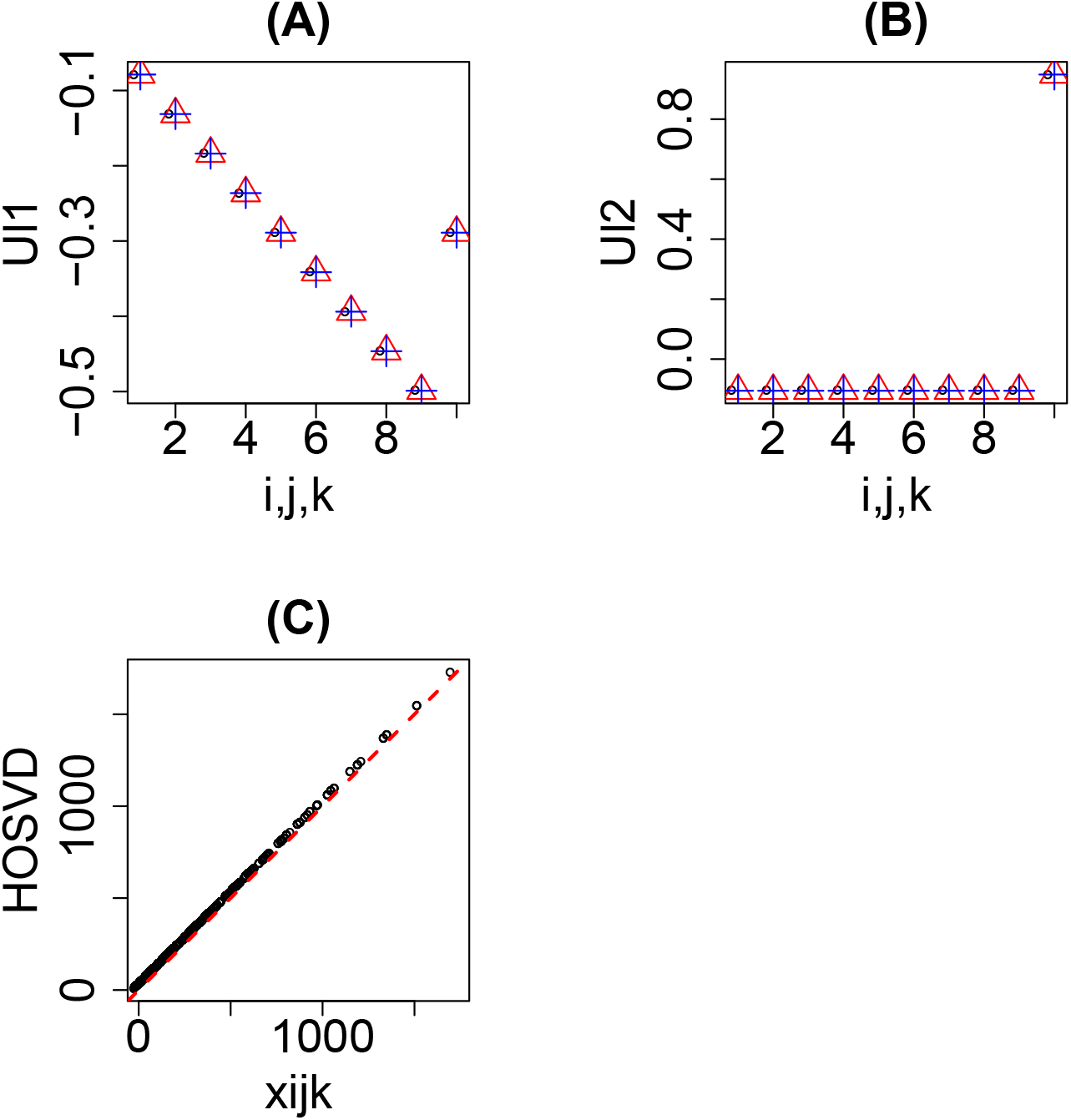
The results obtained by HOSVD applied to data set 2: eq. (5). Other notations are the same as Fig. 2.

It is obvious that HOSVD successfully performs TD (Figs. 2(C) and 3(C)) although obtained singular value vectors (Figs. 2(A) and (B) and 3(A) and (B)) are not equivalent to Fig. 1 because HOSVD assumes the orthogonality between singular value vectors. The first singular value vectors, 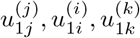 (Figs. 2(A) and 3(A)), clearly represent somewhat means of ***v*** (Figs. 1(A) and 1(C)) and ***v***’ (Figs. 1(B) and 1(D)) while the second singular value vectors, 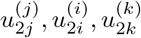 (Figs. 2(B) and 3(B)), clearly represent difference of them.

Next we applied CP decomposition to data set 1 and 2: eqs. (4) and (5) (Fig. 1). It is obvious that CP decomposition (Fig. 4) applied to data set 1 successfully reproduced (Fig. 4(A) and (B)) eq. (3) with eq. (4) (Fig. 1(A) and (B)). On the other hand, CP decomposition (Fig. 5) applied to data set 2 could not, but required up to the third singular value vectors (Fig. 5(A), (B) and (C)). Since CP decomposition depends upon initial values, although we tried multiple initial values, as far as we tried, we could not find the initial values by which CP decomposition can reproduce eq. (3) using eq. (5) (Fig. 1(C) and (D)). In contrast to HOSVD that clearly decomposed ***v*** and ***v***’ into their mean and difference, it is unclear what Fig. 5 represents anymore. Thus, it is obvious whether CP decomposition can perform better than HOSVD is highly dependent upon the data set we analyze. In this sence, HOSVD is less affected by the type of data set analyzed.

**Figure 4:**
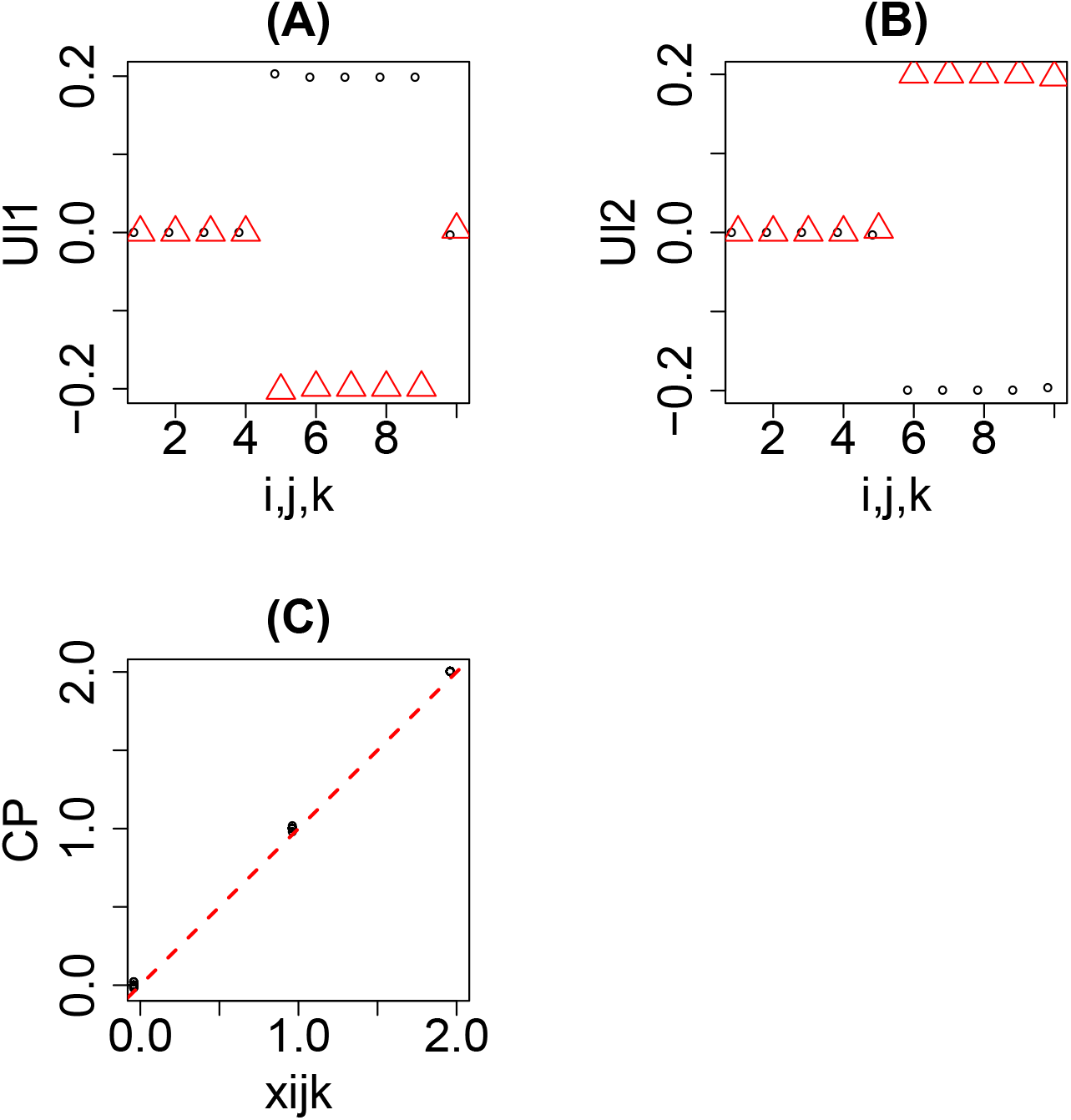
The results obtained by CP decomposition applied to data set 1: eq. (4). (A) Open black circles: 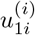, open red triangles: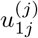, blue pluses: 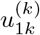 (B) Open black circles: 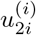, open red triangles: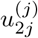, blue pluses: 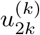. (C) Scatter plot between *x_ijk_* (horizontal axis) and those reproduced by CP decomposition using singular value vectors shown in (A) and (B) (vertical axis).

**Figure 5:**
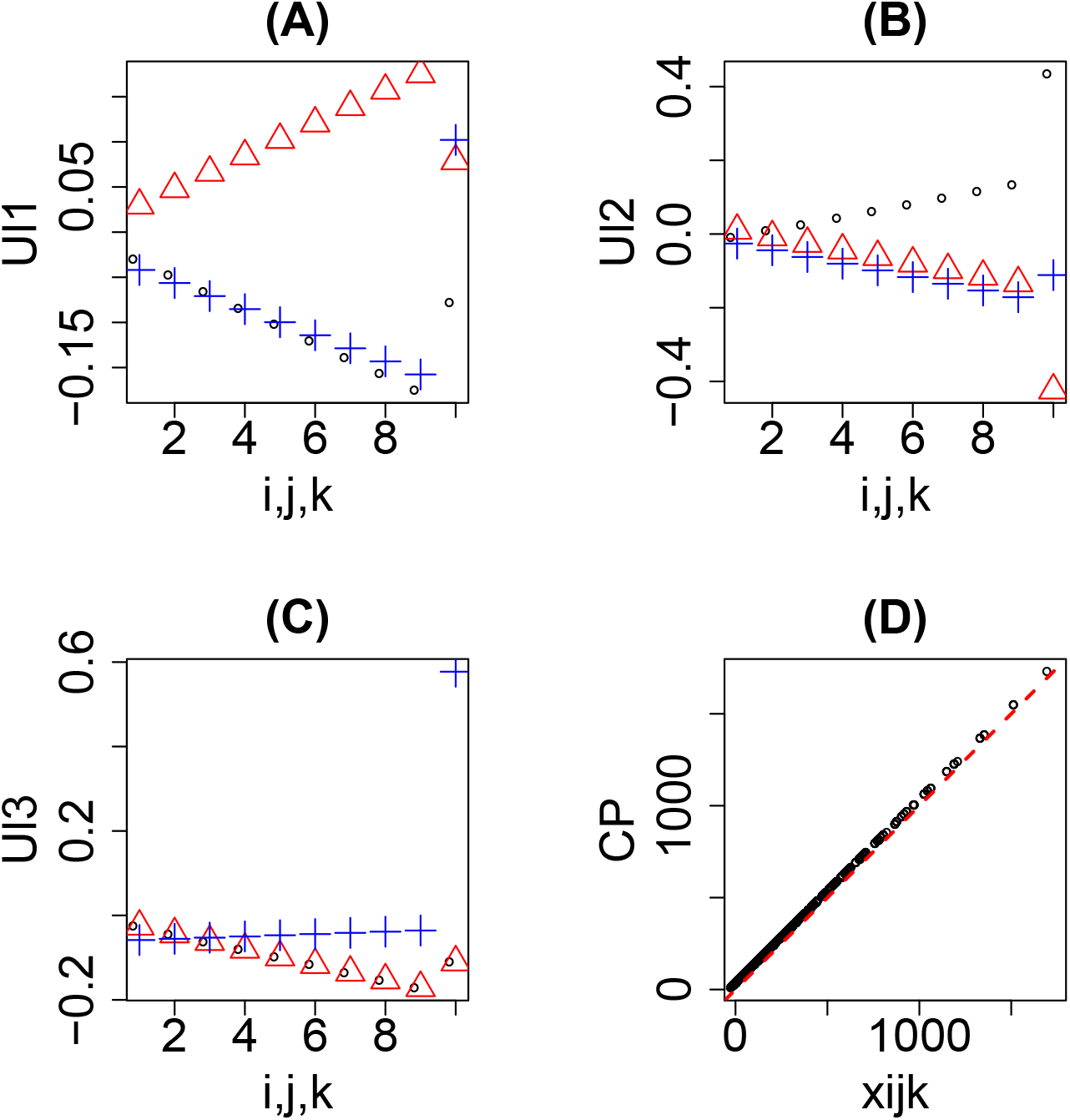
The results obtained by CP decomposition applied to data set 2: eq. (5). (A) Open black circles: 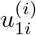, open red triangles: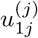, blue pluses: 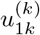 (B) Open black circles: 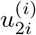, open red triangles: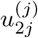, blue pluses: 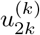. (C) Open black circles: 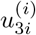, open red triangles: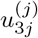, blue pluses: 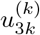. (C) Scatter plot between *x_ijk_* (horizontal axis) and those reproduced by CP decomposition using singular value vectors shown in (A), (B) and (C) (vertical axis).

Finally, we applied CMTF to data sets 1 and 2 (Fig. 1). In order that, we need to specify loss function, *f*, to be minimized;

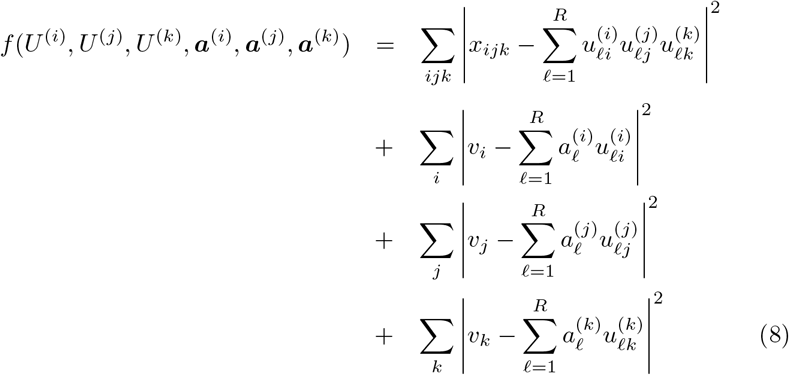

where 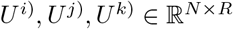 are defined as

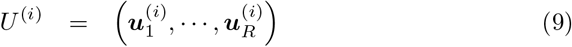

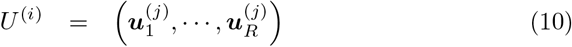

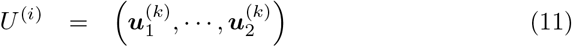

with 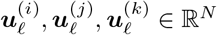 defined as

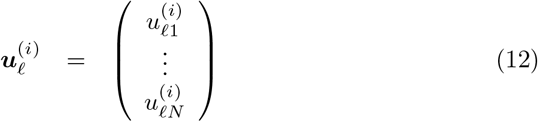

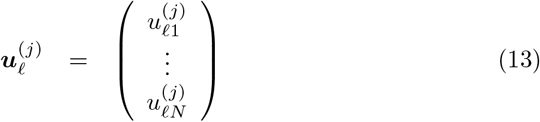

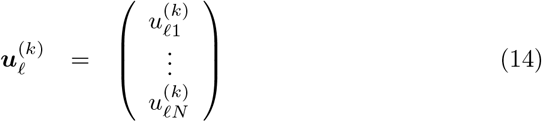

With coefficient vectors, 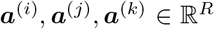, ***v*** is required to be expressed by the linear transformation of *U*^(*i*)^, *U*^(*j*)^, *U*^(*k*)^.

After trying to apply CMTF with *R* =2 (because we know *R* = 2 is enough because of eq. (3)) to data sets 1 and 2, we realized that it is rare that CMTF converges to global minimum when starting from initial values, *U*^(*i*)^, *U*^(*j*)^, *U*^(*k*)^, ***a***^(*i*)^, ***a***^(*j*)^, ***a***^(*k*)^, drawn from 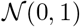 where 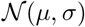 is normal distribution having mean of *μ* and standard deviation of *σ*. After trying several tens of ninital values, we got the results shown in Figs. 6 and 7. It is obvious that CMTF performed quite well as far as it converges. 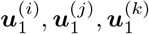, (Figs. 6(A) and 7(A)) correspond to ***v*** (Fig. 1(A) and (C)) while 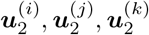 (Figs. 6(B) and 7(B)), correspond to ***v***’ (Fig. 1(B) and (D)) as expected. On the other hand, it is problematic that CMTF rarely converges to global minimum. In order to improve this points, we replaced ALS employed in CMTF with BFGS. Now CMTF came to converge to global minimum (Figs. 8 and 9) with starting any initial values drawn from 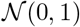 as long as we tried. Thus, we decided to apply CMTF with replacing ALS with BFGS.

**Figure 6:**
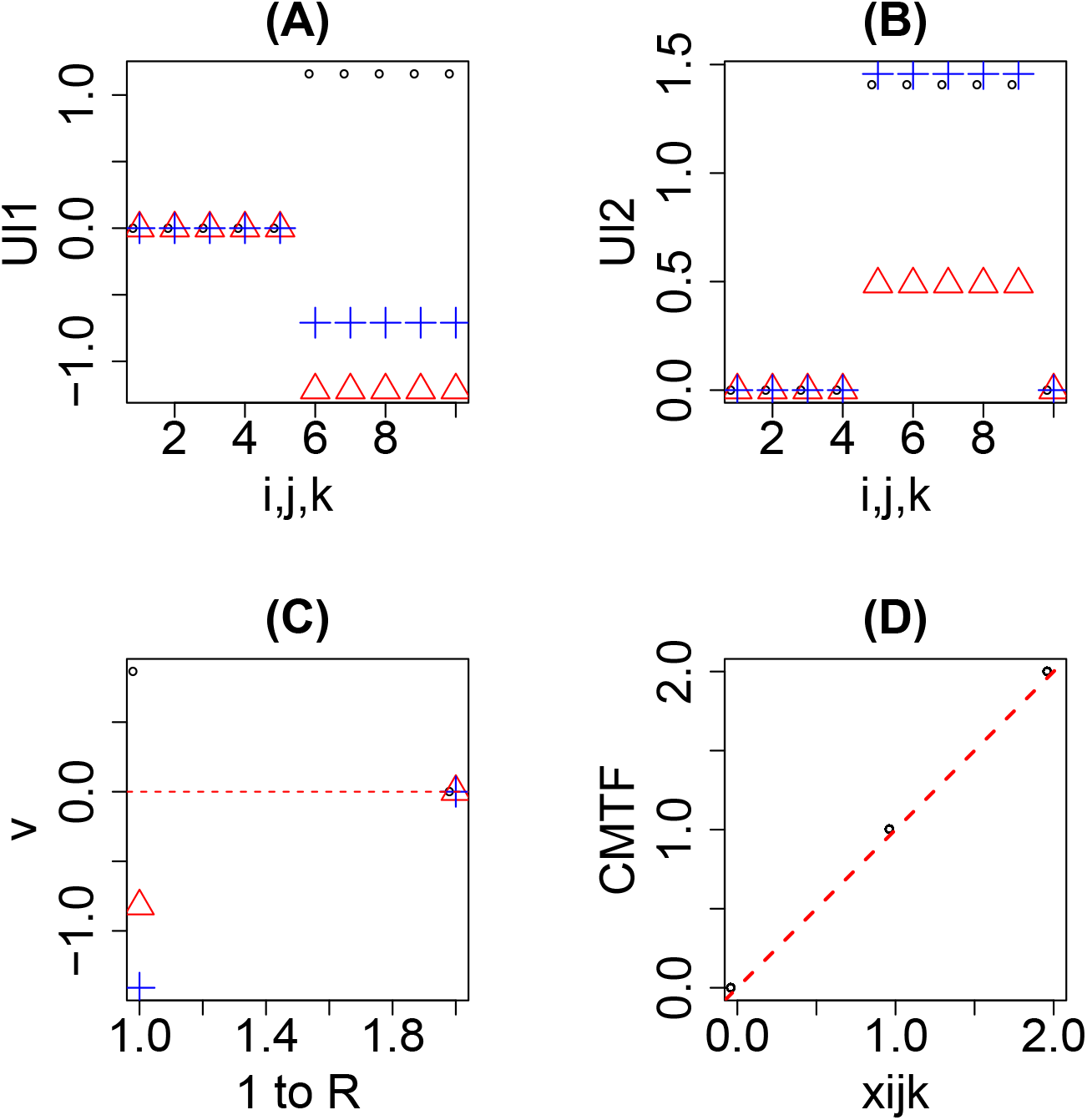
The results obtained by CMTF applied to data set 1: eq. (4). (A) Open black circles: 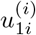, open red triangles: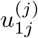, blue pluses: 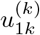 (B) Open black circles: 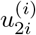, open red triangles: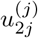, blue pluses: 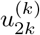. (C) Open black circles: 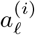, open red triangles: 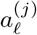, blue pluses: 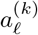. (D) Scatter plot between *x_ijk_* (horizontal axis) and those reproduced by CMTF decomposition using singular value vectors shown in (A) and (B) (vertical axis).

**Figure 7:**
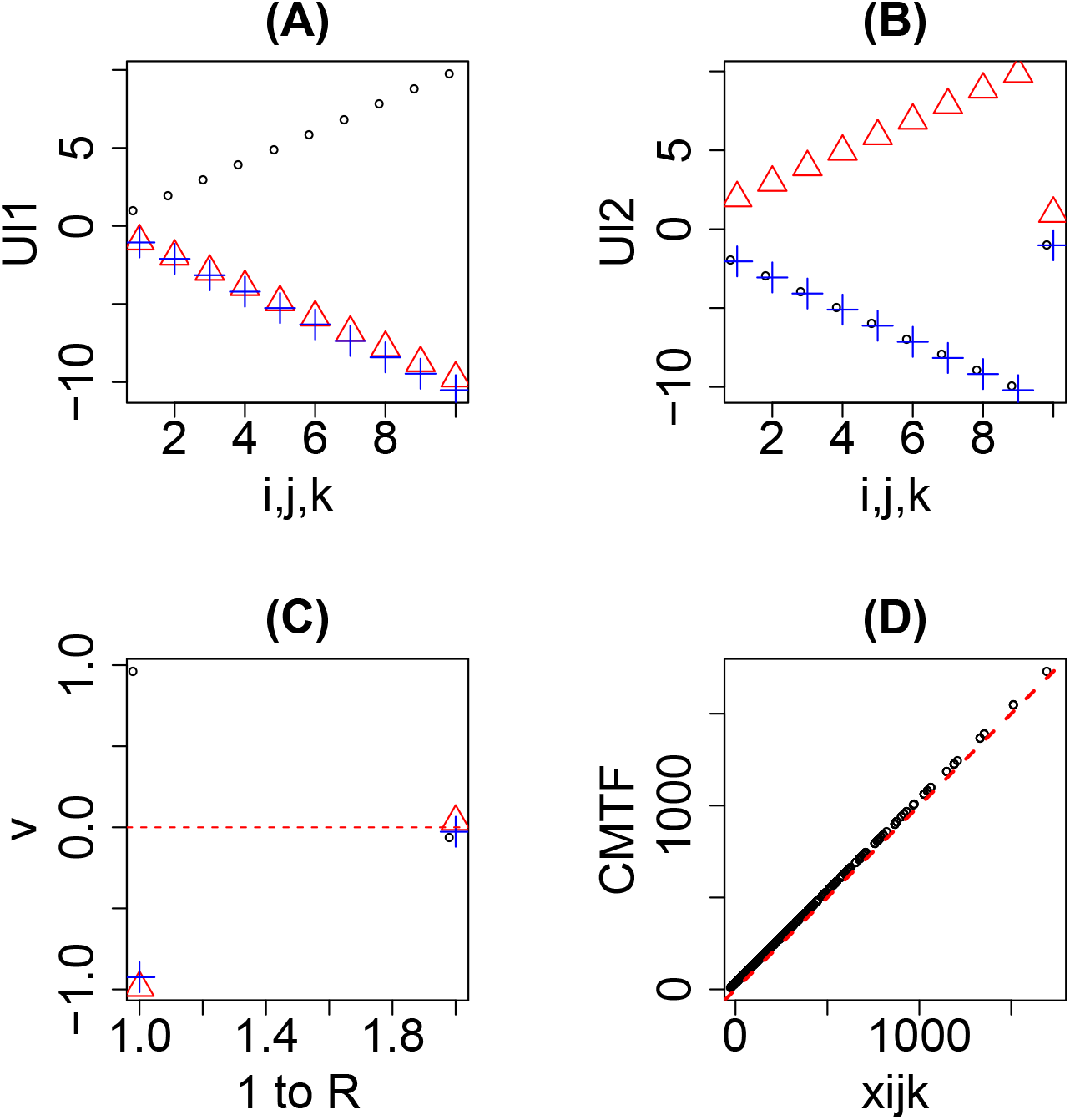
The results obtained by CMTF applied to data set 2: eq. (5). (A) Open black circles: 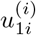, open red triangles: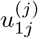, blue pluses: 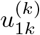 (B) Open black circles: 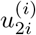, open red triangles: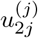, blue pluses: 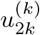. (C) Open black circles: 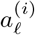, open red triangles: 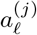, blue pluses: 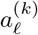. (D) Scatter plot between *x_ijk_* (horizontal axis) and those reproduced by CMTF decomposition using singular value vectors shown in (A) and (B) (vertical axis).

**Figure 8:**
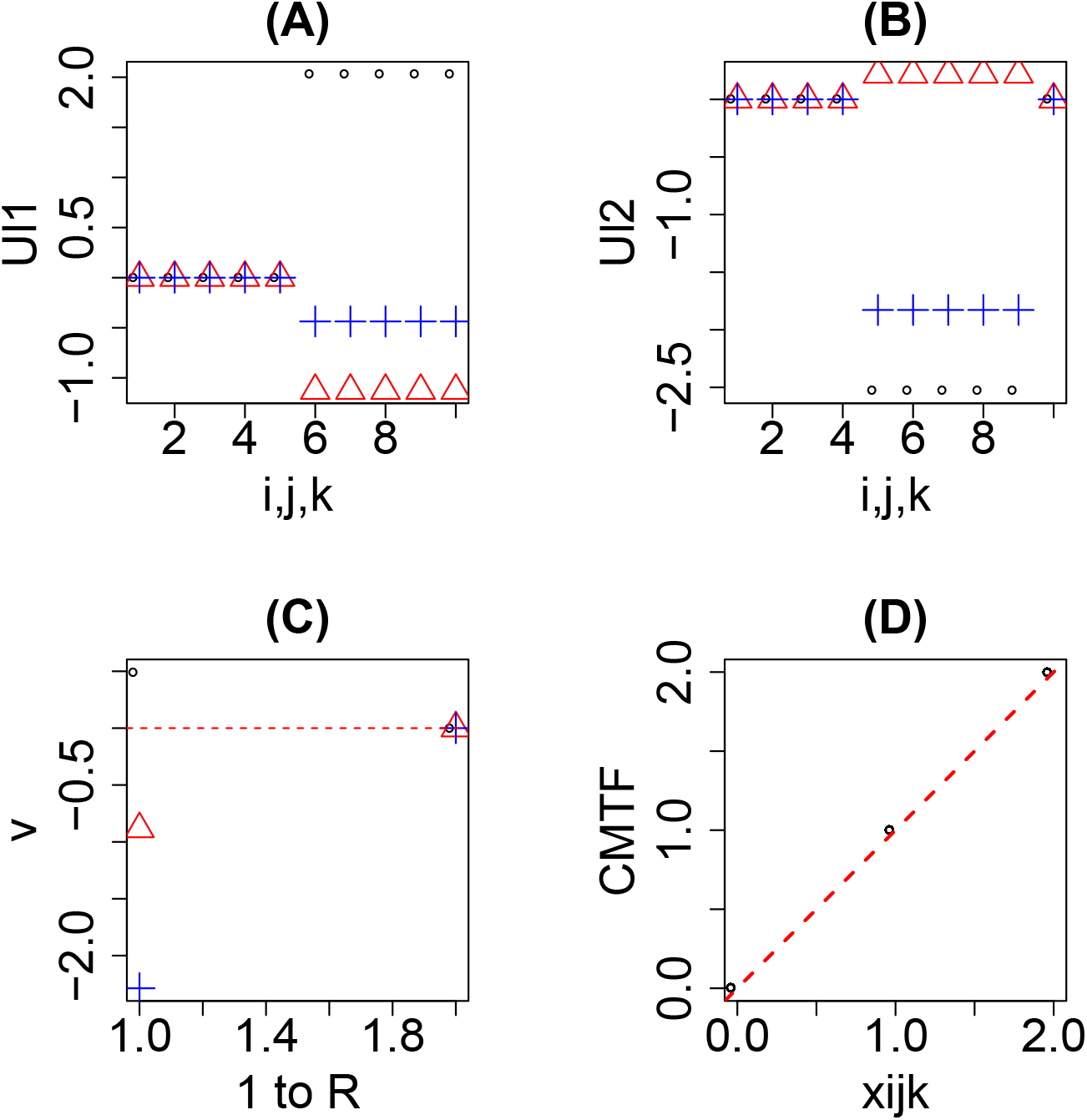
The results obtained by CMTF, with replacing ALS with BFGS, applied to data set 1: eq. (4). (A) Open black circles: 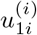, open red triangles: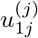, blue pluses: 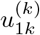 (B) Open black circles: 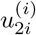, open red triangles: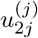, blue pluses: 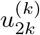. (C) Open black circles: 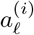, open red triangles: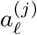, blue pluses: 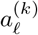. (D) Scatter plot between *x_ijk_* (horizontal axis) and those reproduced by CMTF using singular value vectors shown in (A) and (B) (vertical axis).

**Figure 9:**
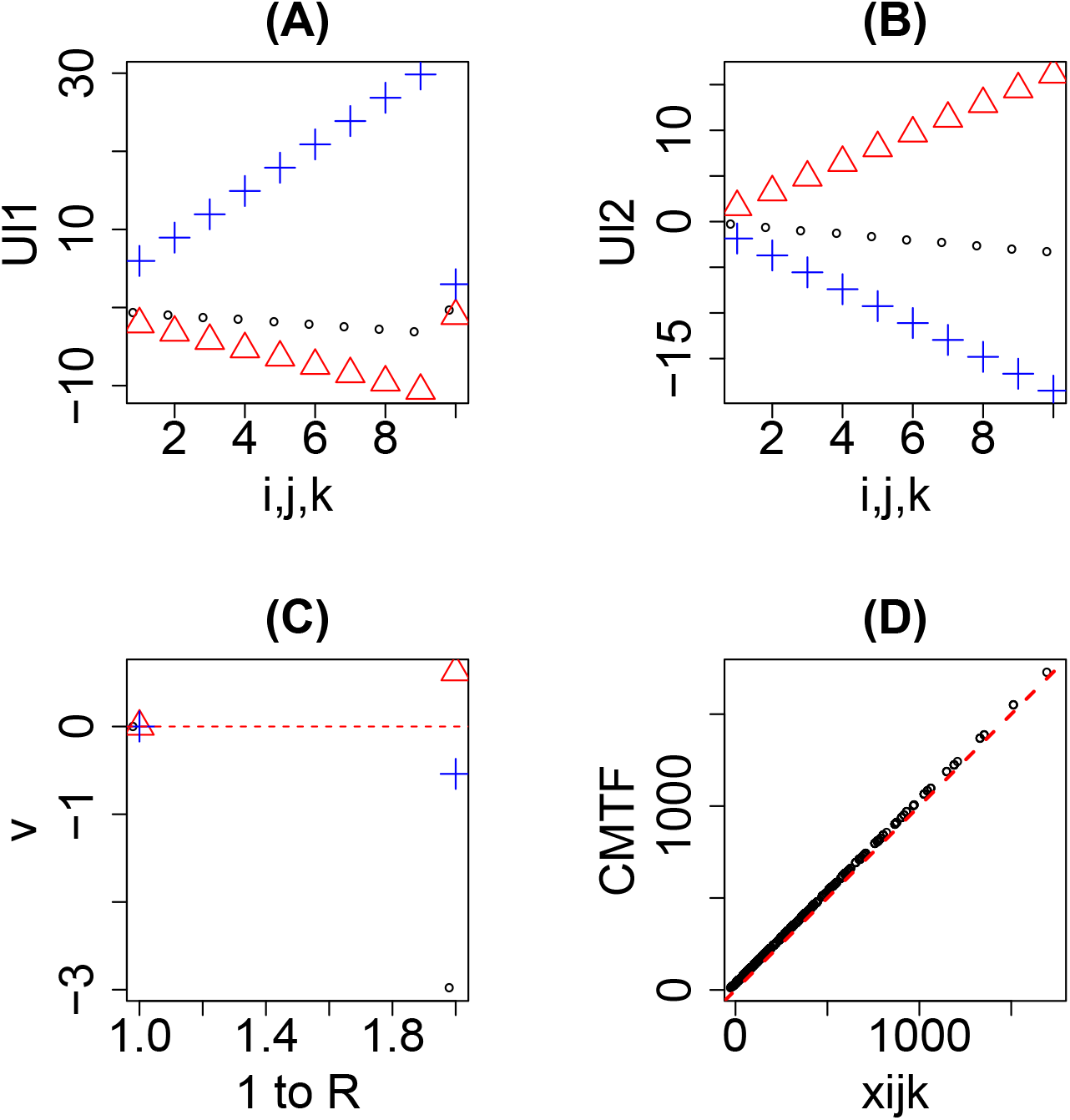
The results obtained by CMTF, with replacing ALS with BFGS, applied to data set 2: eq. (5). (A) Open black circles: 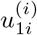, open red triangles: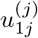, blue pluses: 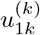 (B) Open black circles: 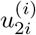, open red triangles: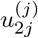, blue pluses: 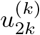. (C) Open black circles: 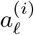, open red triangles: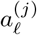, blue pluses: 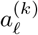. (D) Scatter plot between *x_ijk_* (horizontal axis) and those reproduced by CMTF using singular value vectors shown in (A) and (B) (vertical axis).

Although CMTF looks the best method to apply, CMTF has one problem: cpu time required to perform CMTF. Table 1 shows the list of cpu time required when various metthods are applied to data set 1 and 2. It is obvious that HOSVD is the fastest since it does not require any iterations. CP decomposition is a bit slower than HOSVD, since it requires ALS to converge. CMTF is much more slower no matter which methods, ALS ot BFGS, are employed for the minimization. As far as we deal with small data set, this difference is not critical. Nevertheless, when we have to deal with massive data set, this difference is critical. Although CMTF is slower than HOSVD by only several hundreds times, this difference is generally enhanced when the data set becomes larger. Since cpu time required for HOSVD also increases as data set grows, it might be unrealistic to perform CMTF for much larger data set.

**Table 1:** Cpu time (msec) required to perform various methods.

|  | HOSVD | CP | CMTF |  |
| --- | --- | --- | --- | --- |
|  |  |  | ALS | BFGS |
| data set 1 | 22 | 334 | 5760 | 2002 |
| data set 2 | 9 | 123 | 5787 | 2991 |

Before applying TDs to real data set, we summarize the results here.

- HOSVD is the fastest and its outcome is not affected by the type pf data set much. Nevertheless, because of requirement of orthogonality, it has less ability to derive the structure of original data set, eq. (3), if the vectors used to generate tensor are not orthogonal to each other.
- CP decomposition is the second fastest method and can reproduce the structure of original data set, eq. (3) (Fig. 4). Nonetheless, CP decomposition might fail dependent upon data set (Fig. 5).
- The original CMTF can successfully reproduce the data structure, eq. (3). On the other hand, it is the slowest method and requires to search initial values that converges to global minimum.
- With replacing ALS with BFGS, CMTF comes to converge to global minimum independent of initial values. In spite of the acceleration with this replacement, CMTF is still much slower than HOSVD as well as CP decomposition.

Based upon the observation in the above, since data set we have to analyze is massive, considering primarily the cpu time required, we decided to employ HOSVD first. Then we will try other methods only when HOSVD fails to get reasonable results.

In order to apply the methods to more realistic cases, we added noise to *x_ijk_*. According to the results in the Supplementary file, the summary is as follows:

- HOSVD is least affected by adding noise (Figs. S1 and S2). This is because of the following reason. HOSVD generated two ***u***_*ℓ*_s (Fig. 2(A) and (B), 3(A) and (B)), which correspond to those with larger and smaller amplitudes, respectively, because of the requirement of orthogonality. Then ***u***_*ℓ*_s with larger amplitude remained unchanged (Figs. S1(A) and S2(A)). As a result, correspondence between *x_ijk_* and the reconstruction (Figs. S1(C) and S2(C)) remained relatively accurate.
- For CP decomposition, adding noise destroyed the tiny difference among ***u***_*ℓ*_s (Fig. 4 (A) and (B), Fig. 5 (A), (B) and (C)). Then the CP decomposition could detect only one valid ***u***_*ℓ*_ (Figs. S3(A) and S4(B)). As a result, the obtained ***u***_*ℓ*_ do not look better than those obtained by HOSVD (Figs. S1(A) and S2(A)). Then advantages of CP decomposition over HOSVD, which exist when noise free data set is considered, were lost.
- Original CMTF failed to converge, since adding noise disrupted computation of gradient that is required to update the ***u***_*ℓ*_ by ALS.
- Although CMTF with replacing ALS with BFGS still converged (Figs. S5 and S6), it was impossible to see which ***u***_*ℓ*_ converged correctly, because the converged solution has residuals due to adding noises. As a result, the converged ***u***_*ℓ*_ (Figs, S5(A) and S6(B)) do not look better than those for HODVD (Figs. S1(A) and S2(A)). The correspondence between *x_ijk_* and the reconstruction (Figs. S5(D) and S6(D)) even became worst among methods tested. The advantages over HOSVD, which exist when noise free data set is considered, were lost as for CP decomposition.

In conclusion, adding noise, which is supposed to be closer to a realistic situation, added more advantages to HOSVD than other methods.

### 3.2 Application of HOSVD to real data set

Among numerous neurodegenerative diseases, we focus on Alzheimer’s disease (AD) in this study, because it is the diseases for which the most number of drugs were tried to develop. For example, among 322 drugs that target neurodegenerative diseases, as many as 92 drugs targeted AD [24]. The therapy targets of AD are wide ranged; especially, Amyloid protein was most frequent target (12 among 92 drugs target amyloid), because accumulation of amyloid has ever been believed to be a primary cause of AD.

For this purpose, we selected one specific scRNA-seq data set, GSE127891, by which we can demonstrate the effectiveness of our proposed method. When selecting genes using TD based unsupervised FE, we first need to specify what kind of properties of gene expression we consider. In this study, we require the followings.

1. Gene expression should be independent of cells within the same 96 wells plate.
2. Gene expression should be independent of genotype.
3. Gene expression should be independent of tissues.
4. Gene expression should have monotonic dependence upon age.
5. Gene expression should be independent of sex.
6. Gene expression should be independent of each of four 96 wells plates under the same conditions.

In other words, we try to select genes with the most robust monotonic age dependence as much as possible. The reason of this motivation is as follows. In the paper where data set analyzed here was investigated originally, Frigerio et al. [25] found that age is the primary factor of the microglia response to accumulation of A*β* plaques. We found that singular value vectors with *ℓ*_1_ = *ℓ*_2_ = *ℓ*_3_ = *ℓ*_5_ = *ℓ*_6_ = 1 represent independence of cells, genotypes, tissues, sex and plates (Figure 10 (A), (B), (C), (E), (F)). On the other hand, *u*_2_*j*_4___ represents monotonic dependence upon ages, 1 ≤ *j*_4_ ≤ 4 (Figure 10 (D)).

**Figure 10:**
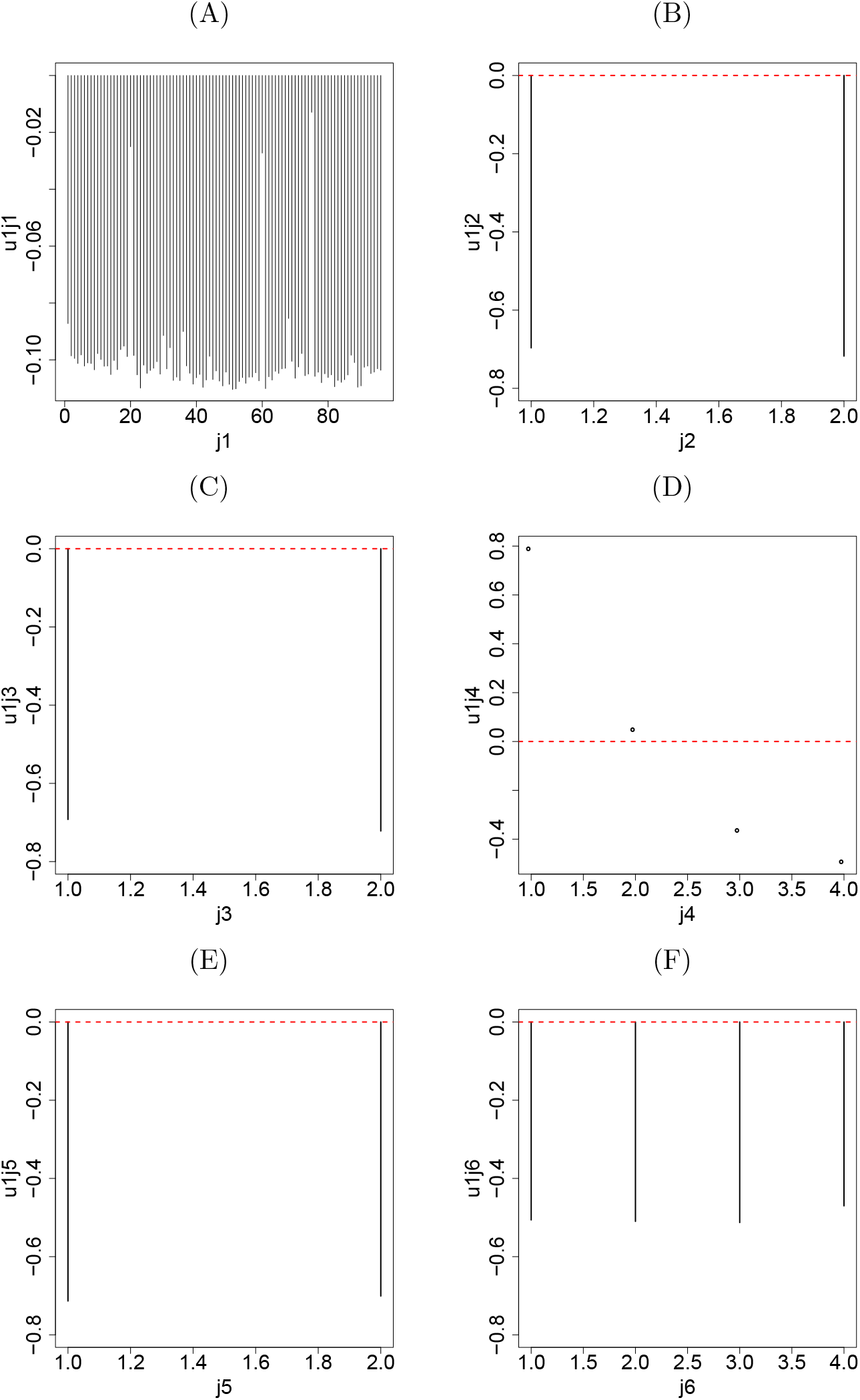
Singular value vectors. (A) *u*_1_*j*_1___ (B) *u*_1_*j*_2___ (C) *u*_1_*j*_3___ (D) *u*_2_*j*_4___ (E) *u*_1_*j*_5___ (F) *u*_1_*j*_6___.

Next, we need to find the *G*(1, 1, 1, 2, 1, 1, *ℓ*_7_) with the largest absolute value in order to identify singular value vector, *u*_*ℓ*_7_*i*_, attributed to genes. Then we found that *G*(1, 1, 1, 2, 1, 1, 2) has the largest absolute value. Therefore, we decided to use *u*_2*i*_ for attributing *P*-values to genes as shown in eq. (2). Finally, 401 genes are identified as being associated with adjusted *P*-values less than 0.01 (The list of genes is available as supplementary material).

These 401 genes are uploaded to Enrichr to identify the compounds, with which genes expressing differential expression of cell lines treated are maximally overlapped with these 401 genes. As for “LINCS L1000 Chem Pert up” category (Table 2, full list is available as supplementary material), the top ranked compound is alvocidib, which was previously tested for AD [26]; there are also 65 experiments (see supplementary material) of cell lines treated with alvocidib and associated with adjusted *P*-value less than 0.05. The second top ranked compound is AZD-8055, which was also previously tested for AD [27]; there are also 6 experiments (see supplementary material) of cell lines treated with AZD-8055 and associated with adjusted *P*-value less than 0.05.

**Table 2:**
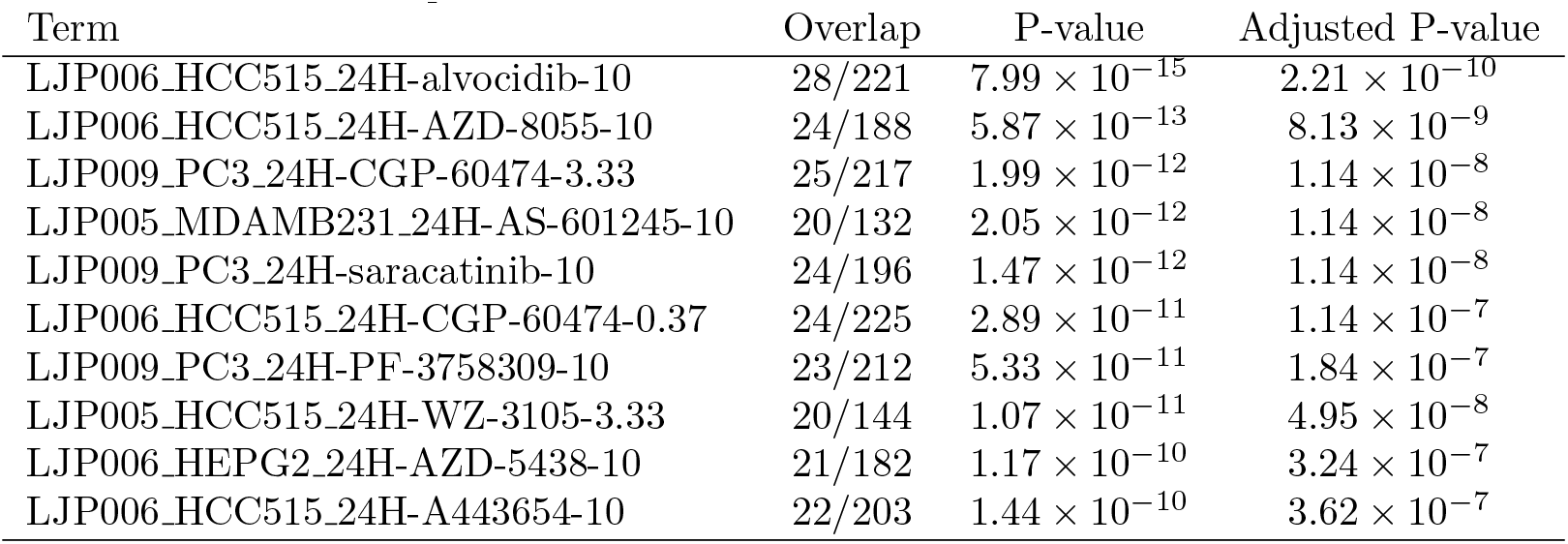
Top ranked 10 compounds listed in “LINCS L1000 Chem Pert up” category in Enrichr. Overlap is that between selected 401 genes and genes selected in individual experiments.

One might wonder if this is an accidental agreement which is specific to LINCS data set. In order to confirm that it is not an accidental agreement, we also see DrugMatrix category (Table 3, full list is available as supplementary material). The top, fifth and tenth ranked compound is cyclosporin-A, which was also previously tested for AD [28];there are also 57 experiments (see supplementary material) of cell lines treated with cyclosporin-A and associated with adjusted *P*-value less than 0.05. Finally, we tested “Drug Perturbations from GEO up” category in Enrichr (Table 4, full list is available as supplementary material). The top ranked compounds is imatinib, which was also previously tested for AD [29];there are also 18 experiments (see supplementary material) of cell lines treated with imatinib and associated with adjusted *P*-value less than 0.05.

**Table 3:**
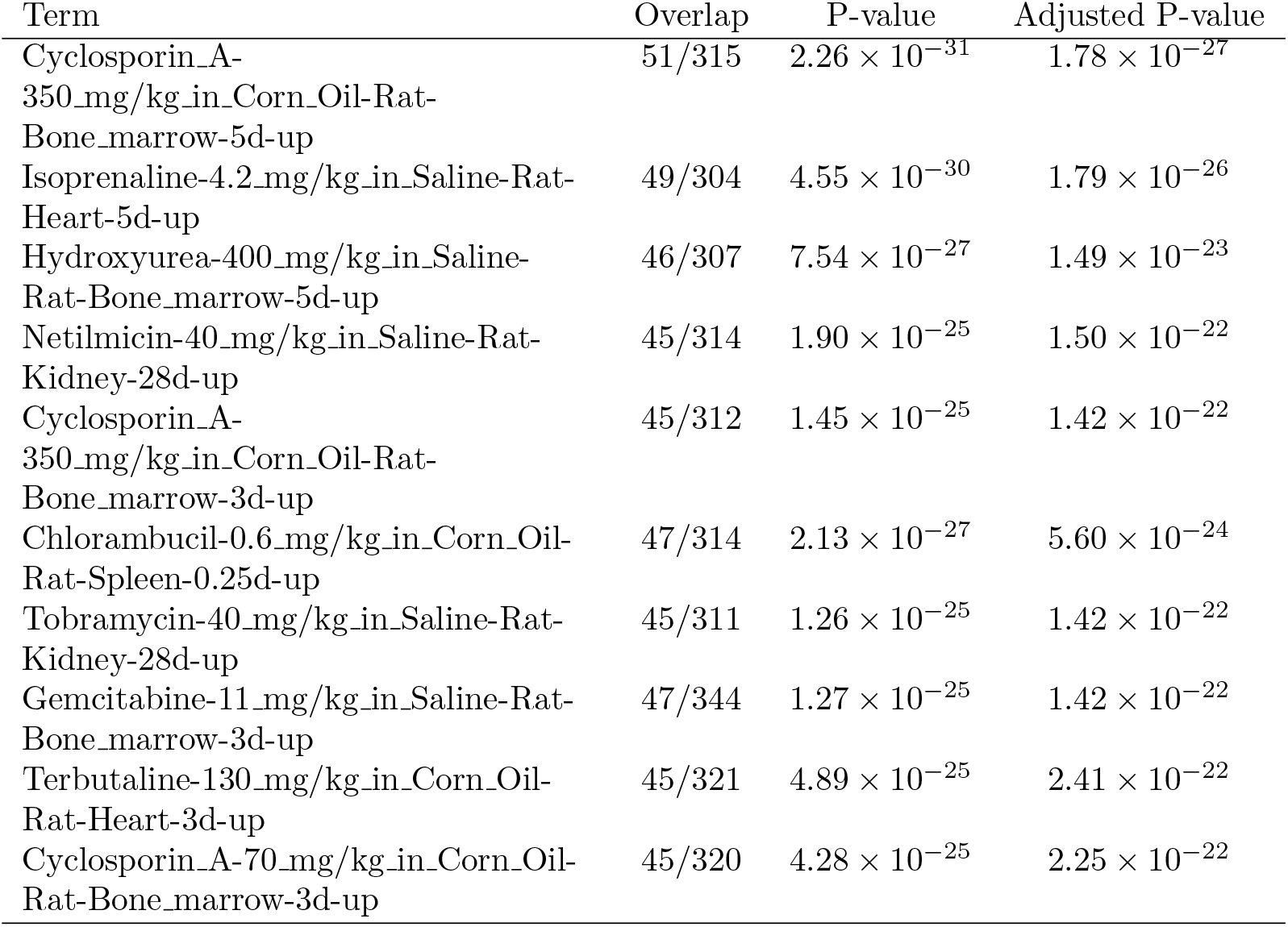
Top ranked 10 compounds listed in “DrugMatrix” category in Enrichr. Overlap is that between selected 401 genes and genes selected in individual experiments.

**Table 4:** Top ranked 10 compounds listed in “Drug Perturbations from GEO up” category in Enrichr. Overlap is that between selected 401 genes and genes selected in individual experiments.

| Term | Overlap | P-value | Adjusted P-value |
| --- | --- | --- | --- |
| imatinib DB00619 mouse GSE51698 sample 2522 | 81/288 | $2.27 \times 10^{-70}$ | $2.05 \times 10^{-67}$ |
| bleomycin DB00290 mouse GSE2640 sample 2851 | 80/329 | $6.09 \times 10^{-64}$ | $2.75 \times 10^{-61}$ |
| soman 7305 rat GSE13428 sample 2640 | 86/532 | $3.87 \times 10^{-53}$ | $3.50 \times 10^{-51}$ |
| coenzyme Q10 5281915 mouse GSE15129 sample 3464 | 76/302 | $6.84 \times 10^{-62}$ | $2.06 \times 10^{-59}$ |
| N-METHYLFORMAMIDE 31254 rat GSE5509 sample 3570 | 70/283 | $2.39 \times 10^{-56}$ | $3.60 \times 10^{-54}$ |
| Calcitonin 16132288 mouse GSE60761 sample 3446 | 65/220 | $8.51 \times 10^{-58}$ | $1.92 \times 10^{-55}$ |
| cyclophosphamide 2907 mouse GSE2254 sample 3626 | 78/413 | $2.47 \times 10^{-53}$ | $2.48 \times 10^{-51}$ |
| Calcitonin 16132288 mouse GSE60761 sample 3447 | 59/177 | $5.88 \times 10^{-56}$ | $7.59 \times 10^{-54}$ |
| PRISTANE 15979 mouse GSE17297 sample 3229 | 71/291 | $1.03 \times 10^{-56}$ | $1.87 \times 10^{-54}$ |
| coenzyme Q10 5281915 mouse GSE15129 sample 3456 | 76/396 | $1.79 \times 10^{-52}$ | $1.35 \times 10^{-50}$ |

In order to check if the results are relatively independent of threshold adjusted P-value, we also checked two additional threshold P-values, 0.005 and 0.05 (See Table 5). Although the threshold adjusted P-values less than 0.01 is the best, other two choices achieve almost similar performance. Thus, the performance achieved seems to be robust.

**Table 5:**
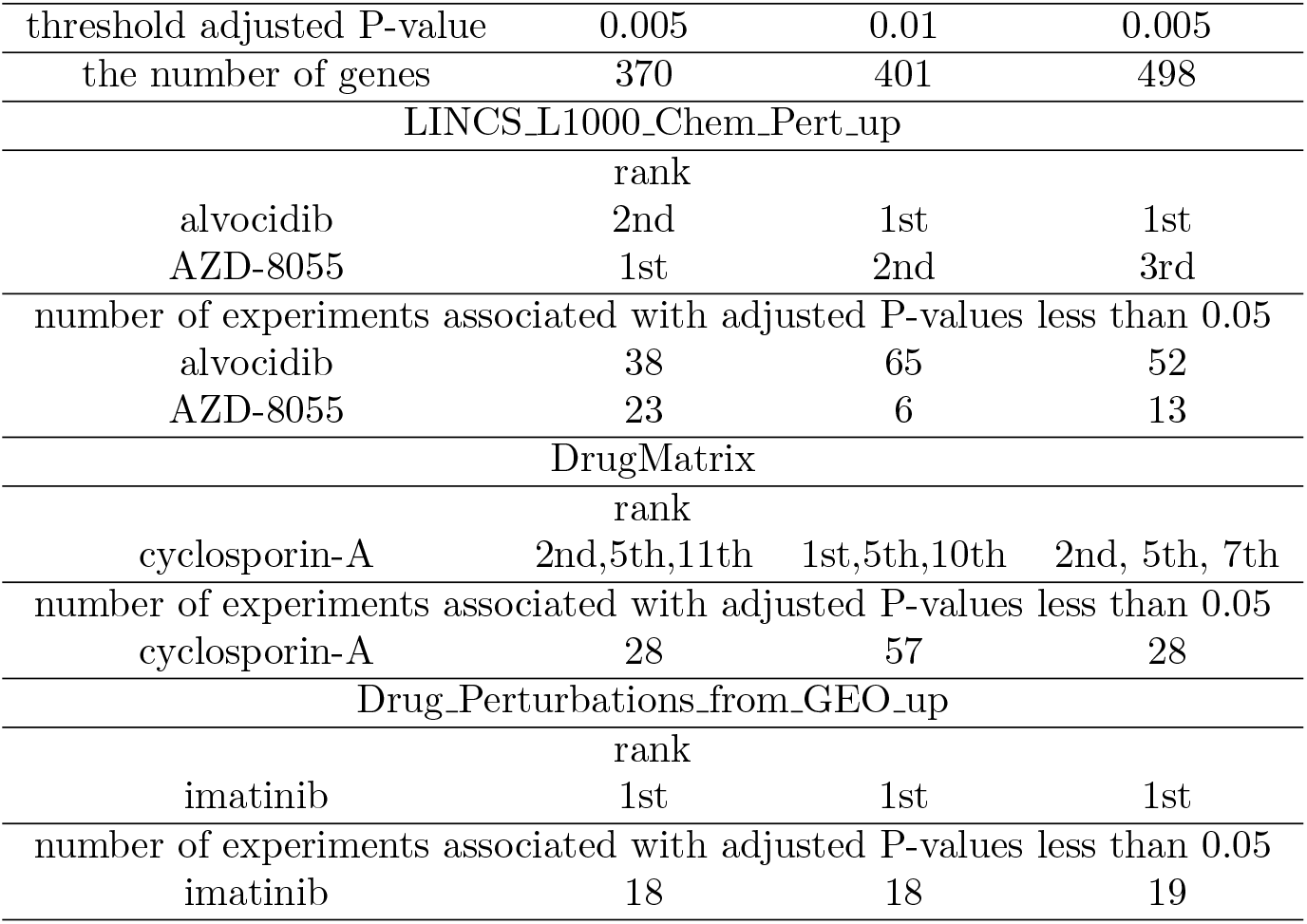
Summary of enrichment analysis for three threshold adjusted P-value

| threshold adjusted P-value | 0.005 | 0.01 | 0.005 |
| --- | --- | --- | --- |
| the number of genes | 370 | 401 | 498 |
| LINCS_L1000_Chem_Pert_up |  |  |  |
|  | rank |  |  |
| alvocidib | 2nd | 1st | 1st |
| AZD-8055 | 1st | 2nd | 3rd |
| number of experiments associated with adjusted P-values less than 0.05 |  |  |  |
| alvocidib | 38 | 65 | 52 |
| AZD-8055 | 23 | 6 | 13 |
| DrugMatrix |  |  |  |
|  | rank |  |  |
| cyclosporin-A | 2nd,5th,11th | 1st,5th,10th | 2nd, 5th, 7th |
| number of experiments associated with adjusted P-values less than 0.05 |  |  |  |
| cyclosporin-A | 28 | 57 | 28 |
| Drug_Perturbations_from_GEO_up |  |  |  |
|  | rank |  |  |
| imatinib | 1st | 1st | 1st |
| number of experiments associated with adjusted P-values less than 0.05 |  |  |  |
| imatinib | 18 | 18 | 19 |

Although these findings suggest that our strategy is effective to find compounds that can be used for AD treatment, one might think that these findings are still weak. Since these 401 genes are simply genes whose expression is altered because of Amyloid accumulation, they themselves are unlikely to be diseas causing genes. Thus we consider regulation factors that affect expression of these genes. At first, we consider transcription factor (TF). With checking “ENCODE and ChEA Consensus TFs from ChIP-X” category in Enrichr, we found that the target genes of TFs, MYC, NELFE, TAF7, KAT2A, SPI1, RELA, TAF1 and PML are top ranked ten TFs associated with adjusted P-values less than 1 × 10^−7^ (They are less than ten, because some are ranked in multiple times within top 10). Among them, MYC [30], KAT2A [31], SPI1 [32], RELA [33], TAF1 [34], and PML [35] were reported to be related to AD. These TFs were also identified within top ranked 10 TFs, with other two additional threshold P-values, less than 0.005 and 0.05, with similar associated adjusted P-values; no additional TFs were ranked within top 10.

Next we consider microRNA (miRNA) as regulatory factors towards identified 401 genes. With checking “miRTarBase 2017” category in Enrichr, we found that target genes of miRNAs, hsa-miR-320a, hsa-miR-1260b, hsa-miR-652-3p, hsa-miR-744-5p, hsa-miR-16-5p, hsa-miR-100-5p, hsa-miR-615-3p, hsa-miR-484, hsa-miR-296-3p, and hsa-miR-423-5p are top ranked ten miRNAs associated with adjusted P-values less than 1 × 10^−3^. Among them, miR-320a [36], miR-652 [37], miR-744 [38], miR-16 [39], miR-100 [40], miR-615 [41], miR-484 [42], miR-296 [43], and miR-423 [36] were reported to be related to AD. As for additional two threshold adjusted *P*-values, all are ranked within top 10 for adjusted P-values less than 0.05 while eight out of ten excluding miR-615-3p and miR-296-3p are ranked within top 10. Thus, it also shows a robust result.

These finding can add more confidence that identified 401 genes are likely related to AD. Expression of these 401 genes might be altered because they are simply downstream genes caused by AD, it is unlikely to find more direct evidence that these genes really contribute to AD directly. For our purpose, screening drugs with gene expression, 401 genes are enough to be downstream genes caused by AD. Thus, we do not investigate biological background of these 401 genes further.

Thus, it might be worthwhile investigating lower ranked compounds in Tables 2, 3 and 4 as candidate compounds for AD, even if they were not known drugs for AD.

## 4 Discussion

First of all, since these cell lines in Table 2 are originated in human, our strategy can provide us the opportunity to check if proposed candidate drugs screened with model animals are also effective in human.

It is also remarkable that we do not need gene expression of all genes, but only a subset of genes (please remember that LINCS project measures only gene expression of less than one thousand genes) in order to predict candidate drugs with high accuracy. This might reduce the amount of money to screen numerous number of compounds.

Our method is also applicable to scRNA-seq in order to screen drug compounds candidate from scRNA-seq. To our knowledge, there are very limited number of studies that relate scRNA-seq to drug design [44,45], since scRNA-seq usually lacks cell labeling which is useful to screen differentially expressed genes. In this study, we simply make use of ages, which is not always directly related to diseases. In spite of that, drug we listed was correct, i.e., they are known drugs to some extent. Therefore, our strategy is also useful to add an alternative one along this direction, i.e., making use of scRNA-seq for drug design.

Thus, our strategy, TD based unsupervised FE, might be promising methodology to screen drug candidate compounds.

One might wonder why we have specifically used HOSVD algorithm although there are many other ways by which we can apply TD to data set. There are multiple reasons why we did not employ other TD based approaches. First of all, we would like to compare HOSVD with other simple (unsupervised) TDs, CP decomposition, HOOI for Tucker decomposition and tensor train decomposition. CP decomposition is the much more popular methods because it can relate singular value vectors one to one. In HOSVD algorithm, we need to investigate core tensor, *G*, for relating sigular value vectors attribted to genes an those attreibuted to individual cells. In CP decomposition, since TD is composed of outer product of individual singular value vectors, it is clear which singular value vectors attributed to genes are associated with selected singular value vetors attributed to cells. Nevertheless, CP decomposition has two disadvantages: massive computational time and the lack of guarantee that converges to unique solutions. Since CP decomposition employed alternative least square (ALS), it needs to initial values of singular value vectors, which often converges to distinct final singular value vectors. This results in distinct set of genes selected, since we make use of singular value vectors attributed to genes in order to select genes. It definitely prevents us from interpreting biological meanings that should be independent of numerical initial values. The employment of ALS also results in the lack of estimated computational time, since it is iterative procedure. Especially when we need to deal with massive data set that require huge cpu time in each iteration, it is not a good strategy to employ the method that requires iterative processes that we cannot estimate the cpu time require by it in advance. On the other hand, HOSVD is essentially SVD of unfolded tensor, thus it does not require any iterative computation; it is guaranteed to converge within polynomial time. Since we could get reasonable results using HOSVD, we have no motivation to employ the method that requires iteration like CP decomposition. As for HOOI, since it also employed ALS, it is not recommended to employ for the massive data set that we analyzed in this study. Especially, since it is very usual that HOOI employs the results of HOSVD as initial (starting) values for the iteration, there are no reasons to apply HOOI to the results of HOSVD that is good enough in this study. Finally, as for tensor train decomposition, it does lack the weight factor that relates between singular value vectors attributed to gene and cells. Since we definitely need to relate them for our purpose, tensor train decomposition is not a suitable method, either. All of these point about the comparisons between HOSVD and other TDs from the point of views of feature selection was discussed in more details in the book [9] to be published soon.

After that, we would like to discuss why we do not employ more advanced supervised methods. In the above analysis, we made use of labeling information, e.g., sex, genotypes, and time points, only after TD was applied to data set. On the other hand, there are multiple methods that can make use of labeling information with applying TD. For example, coupled matrix and tensor factorization (CMTF) [23] is a straight extension of unsupervised TD to supervised one. CMTF requires that linear combination of singular value vectors must be coincident with given labeling attributed to samples (in this study, cells). Although it is generally expected that CMTF can derive singular value vectors that are more associated with labeling than fully unsupervised TDs do, only one obstacle to perform CMTF is cpu time. Since CMTP requires iterative optimization to fullfil the requirements, i.e., linear combination of singular value vectors must be coincident with given labeling attributed to sample, CMTF requires more computational time than unsupervised TD including HOSVD do. Practically, CMTF requires as many as hundreds itetartions, each of which requires cpu time as much as HOSVD requires. This means, CMTF takes as many as hundreds times longer that HOSVD. In this case, since data set is so massive, single HOSVD requires several hours run on computer, Although we tried to implement CMTF fitted to our model and to execute it, it does not converges within a day. Since our TD based unsupervised FE has already achieved reasonable results we concluded that performing more advanced supervised methods that usually require more cputime is not effective and did not employ any supervised method including CMTF.

## 5 Conclusion and Future Work

In this paper, we applied TD based unsupervised FE to scRNA-seq taken from mouse brain with A*β* accumulation. We have compared selected 401 genes with differentially expressed genes in cell lines and model animals treated with various compounds. As a result, as for three independent data sets, LINCS, DrugMatrix and GEO, top ranked compounds are reported to be tested as AD treatment. This suggests the effectiveness of our strategy and lower ranked compounds should be tested as promising drug compounds candidates. To our knowledge, this is the first successful one that can be applied to scRNA-seq in order to identify drug compounds candidate.

For future work, we aim to (1) utilize the tensor decomposition technique in the transfer learning setting to identify effective drugs between target and related tasks in various problems in the clinical informatics domain, among other uses; (2) add other data source of different diseases (e.g., Parkinson’s disease) for treatment validation; and (3) apply the tensor decomposition technique in more fields such as social networks to verify its effectiveness in applications such as recommender systems.

## Supporting information

Gens and enrichment analysis

Supporting information for synthetic data with noises

R code to reproduce the result

## 6 Acknowledgement

This project was funded by the Deanship of Scientific Research (DSR) at King Abdulaziz University, Jeddah, under grant no. KEP-8-611-38. The authors, therefore, acknowledge with thanks DSR for technical and financial support.

This paper was also funded by KAKENHI 19H05270 and 17K00417.

