## Supporting information for synthetic data with noises for "Neurological disorder drug discovery from gene expression with tensor decomposition"

### Supplementary document

In order to test various methods employed in “Synthetic study of TDs” in main text in more realistic situation, we added noise to $x_{ijk}$ as follows.

For data set 1: $x_{ijk} \rightarrow x_{ijk} + \mathcal{N}(0, 1)$

For data set 2: $x_{ijk} \rightarrow x_{ijk} + \mathcal{N}(0, 300)$

$\mu$s are decided such that amplitude should be the same order between signals and noises.

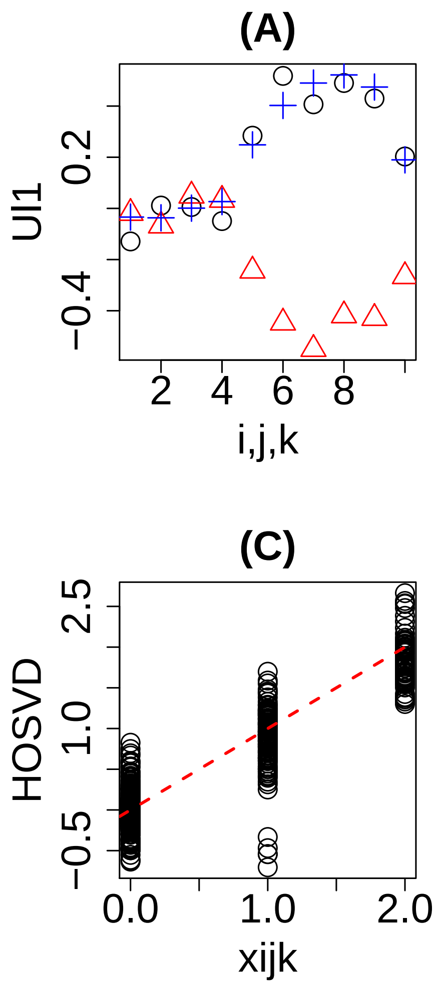


Figure S1: Results of HOSVD applied to data set 1 with adding noise. It corresponds to Fig. 2 in main test

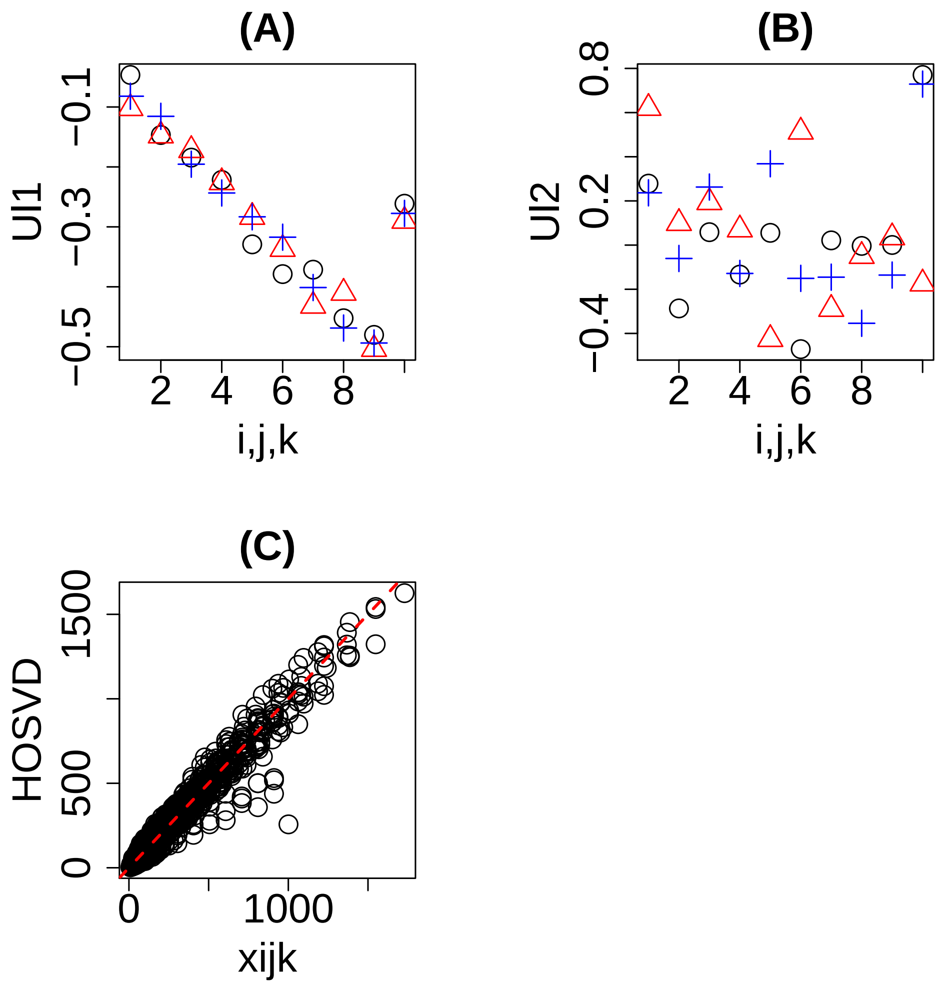


Figure S2: Results of HOSVD applied to data set 2 with adding noise. It corresponds to Fig. 3 in main test

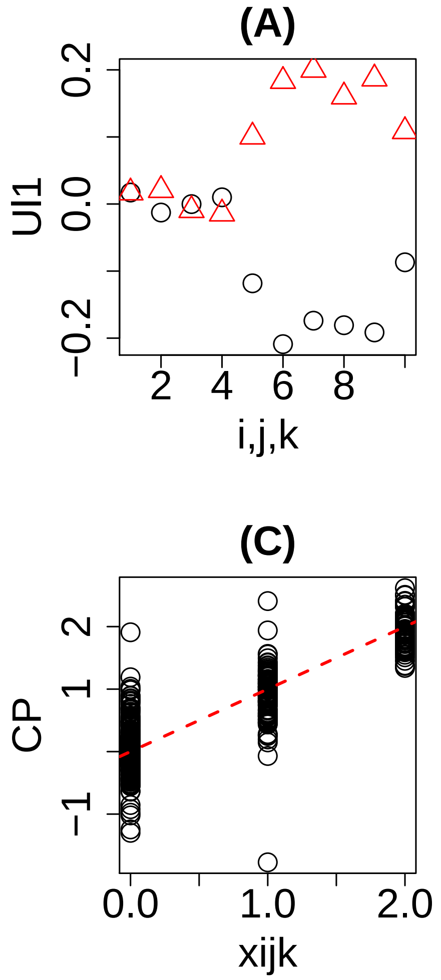


Figure S3: Results of CP decomposition applied to data set 1 with adding noise. It corresponds to Fig. 4 in main test

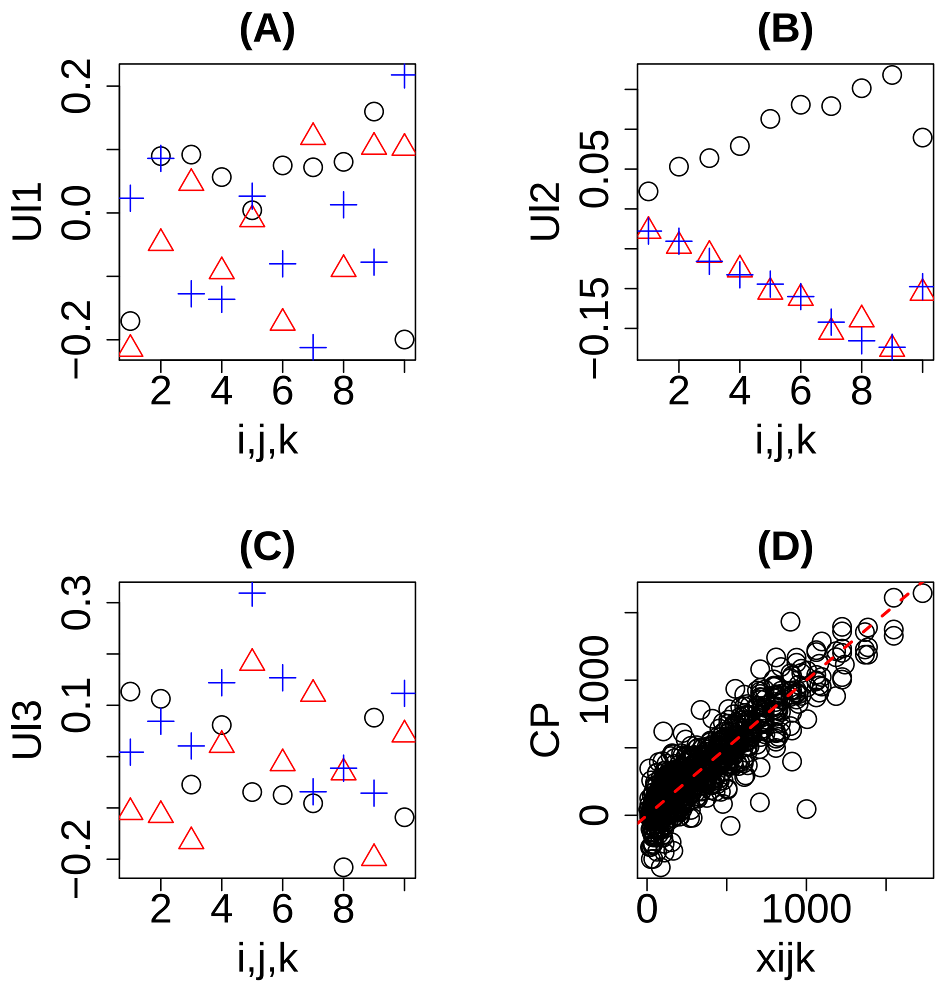


Figure S4: Results of CP decomposition applied to data set 2 with adding noise. It corresponds to Fig. 5 in main test

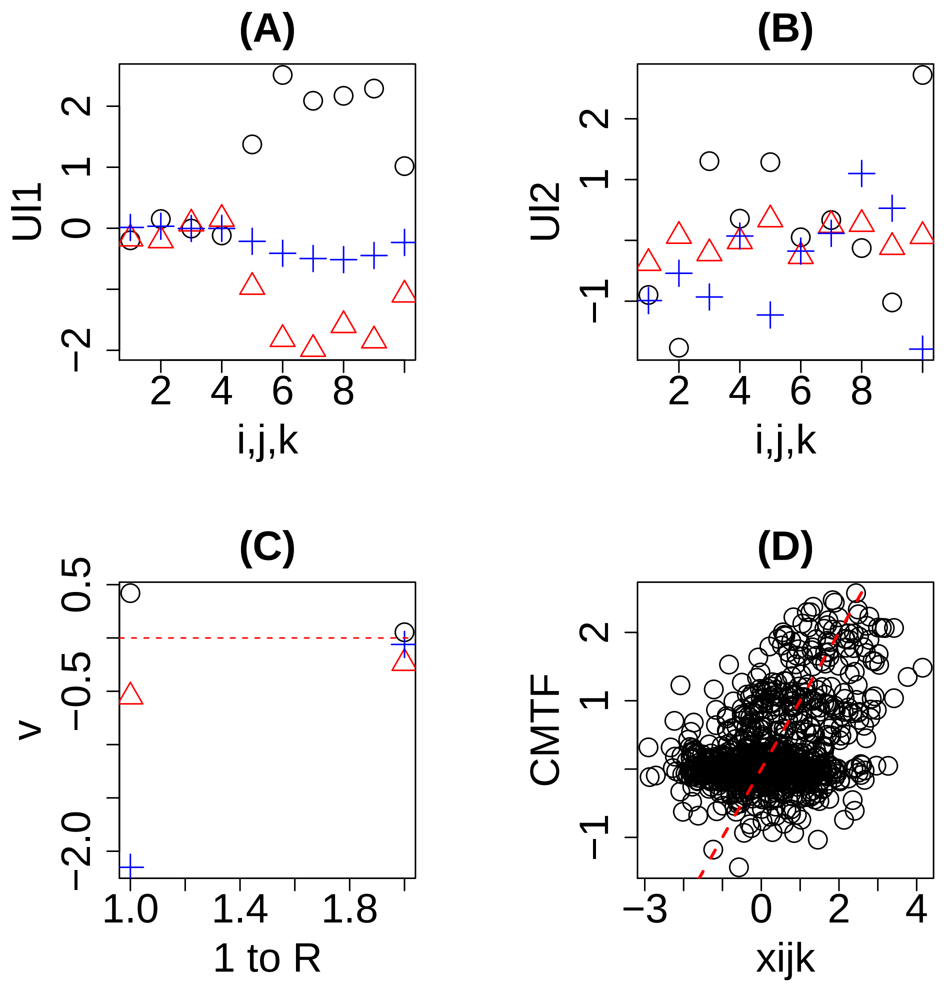


Figure S5: Results of CMTF, with replacing ALS with BFGS, applied to data set 1 with adding noise. It corresponds to Fig. 8 in main test

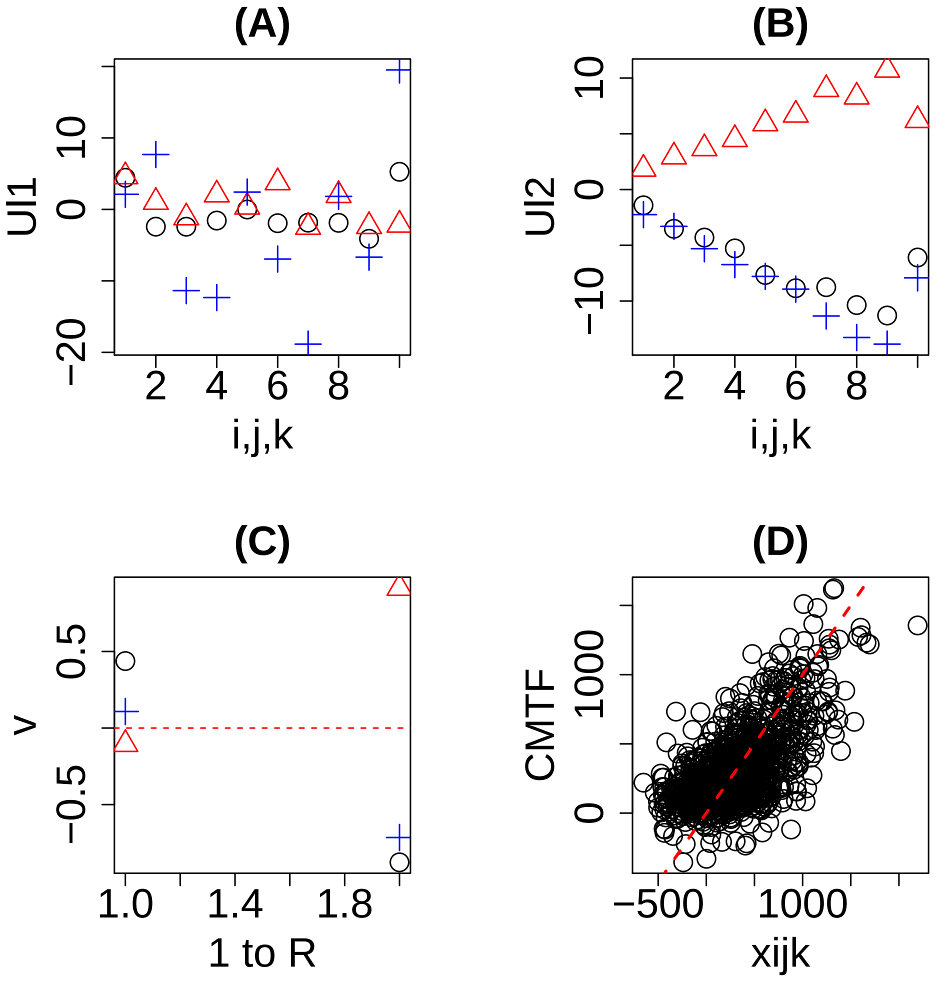


Figure S6: Results of CMTF, with replacing ALS with BFGS, applied to data set 2 with adding noise. It corresponds to Fig. 9 in main test
